# Degree-ranked gene lists omit the cross-module connectors, and a partition-free centrality recovers them

**DOI:** 10.64898/2026.08.10.743862

**Authors:** Zhao Qun, Zheng Huaizheng, Zhang Yuxin, Bi Jieying, Sun Tan

## Abstract

Network centrality is the workhorse of gene prioritisation, yet what a ranking omits is rarely audited. Scoring each selection against an annotation-count-matched maximum-entropy reference—asking whether a selected gene set covers the genome’s functional space or collapses it-reveals that the criterion in standard use has a measurable blind spot in exactly the class it is meant to surface. Degree, the most widely used criterion, returns the cross-module bridges that are also locally dominant—connector *hubs*—and omits the non-hub connectors: where 26% of the genome occupies these coordinating roles, a degree-ranked list holds 18% and an EDVS-ranked list 55%, and degree’s top-1% collapses functional coverage below the reference on all five networks tested. We repurpose EDVS (Entropy of Degree-Vector Sums), an information-theoretic diversity measure, as an annotation-free, partition-free centrality that recovers this omitted class. The coverage it preserves is carried by cross-module participation *P*, which cannot be computed without a community partition; EDVS matches *P*-level coverage on all five networks using none, and retains 0.84 of its selection under edge perturbation that leaves partition-based selections at 0.21-0.46. The deficit is general: the collapse holds in the same direction on the two networks built without functional annotation (0.5–1.1 bit; co-expression, physical interaction) as on the three supervised by it (1.6–3.3 bit; RiceNet, AraNet, STRING), so supervision amplifies it rather than creates it. The remedy is bounded: EDVS ceases to preserve coverage on the sparse physical-interaction network. And the class EDVS isolates is organizational, not an importance signal: pre-registered probes—essentiality, transcription-factor identity, tissue-specificity, date/party-hub character, phenotype co-localisation—return null or reversed throughout. The conclusive ones are equivalent to their degree-matched nulls within ±5 percentage points (demonstrated, not merely undetected), and the classical coupling of centrality to importance itself holds only network-dependently.

**Author Summary:** Genes rarely act alone: many diseases and agricultural traits are shaped by genes that coordinate several biological processes rather than specialising in one. The standard way to find such genes in a network of gene interactions is to count each gene’s connections—its “centrality”—and rank genes by that count. We show this standard approach has a blind spot: it favours genes that dominate one process over genes that quietly bridge several processes without dominating any, and this blind spot appears across rice, thale cress, and yeast gene networks. We repurpose a diversity measure from an unrelated field (originally used to compare citation patterns) as a new way to rank genes that finds these bridging genes from network structure alone, without needing gene-function annotations—which are themselves incomplete and biased toward well-studied genes—or a prior, unstable step of splitting the network into modules. We are careful to show where the new approach also falls short: on sparse, noisy networks it stops working, and the genes it recovers are not shown to be more biologically important than other genes, only differently positioned. What that position is for is a question this work leaves open.

## 1 Introduction

A recurring goal across functional genomics is to identify the genes that *coordinate* across biological processes rather than those that specialise within one—the pleiotropic, cross-cutting genes whose perturbation reverberates through multiple traits [Paaby and Rockman, 2013]. In network terms these are the genes that bridge otherwise separable functional modules, and the dominant strategy for finding them is network centrality: because biological networks are scale-free and their hubs are enriched for essential genes, high-centrality nodes have long been read as the functionally most important [Barabási and Albert, 1999, Albert, 2005, Jeong et al., 2001], a premise built into candidate-gene prioritisation and function-prediction pipelines alike [Moreau and Tranchevent, 2012, Mostafavi et al., 2008]. That premise, however, folds together two properties that need not coincide: a gene’s *importance*, in the essentiality sense, and its *organizational role* as a cross-module connector. The distinction is sharpest exactly where it is least attended—in crop and other non-model species, where the coordinating genes that shape trait architecture are the practical target, yet the tools for finding them are inherited from an importance-centred hub literature.

A second problem compounds the first. The curated knowledge used to judge whether a centrality measure “works”—GO and pathway annotations—is itself heavily biased toward genes that have already been studied, and study effort tracks degree [Gillis and Pavlidis, 2011, Pavlidis and Gillis, 2012, Stoeger et al., 2018, Haynes et al., 2018, Schaefer et al., 2015]. Well-connected genes are better annotated not only because they are important but because they are visible, and a large fraction of the genome remains uncharacterised for reasons closer to sociology than to biology [Kustatscher et al., 2022, Wood et al., 2019, Edwards et al., 2011]; the gap is widest in plants, where most genes still lack experimental annotation [Rhee and Mutwil, 2014, Rhee et al., 2008]. Judging a degree-correlated hub detector by degree-biased curated knowledge is therefore partly circular, and guilt-by-association inference inherits the same bias [Gillis and Pavlidis, 2012]. A measure computed from topology alone breaks this circle—so long as it is then evaluated for what it *characterises* rather than assumed to track importance.

The organizational axis this importance framing overlooks is well described. Guimerà and Amaral separate provincial hubs, locally dominant within one module, from connectors, whose edges span modules [Guimerà and Amaral, 2005]; the date/party distinction [Han et al., 2004] and the recurrence of connector hubs across biological and neural networks [van den Heuvel and Sporns, 2013] point to the same category. Capturing it, however, has been costly. The participation coefficient that formalises the connector role requires a community partition, inheriting the instabilities and resolution-dependence of community detection [Fortunato, 2010, Fortunato and Barthélemy, 2007, Traag et al., 2019]. Degree, and Burt’s structural-hole constraint [Burt, 1992, 2004] —which we find empirically degenerates onto degree in these dense networks—index importance rather than bridging. Betweenness captures bridging without a partition, but at the *O*(*nE*) cost of all-pairs shortest paths [Brandes, 2001], with sensitivity to network noise [Borgatti et al., 2006], and, as we show, in dense co-functional networks it returns bridging nodes that remain locally central—higher-*z* connector hubs rather than the purely cross-module connector.

Entropy-based centralities are natural candidates for measuring diversity of connection [Fei and Deng, 2017, Qiao et al., 2017, Qiu et al., 2021], yet they, and the broader vital-node and influential-spreader literature [Chen et al., 2012, Kitsak et al., 2010, Lü et al., 2016, Opsahl et al., 2010], have been evaluated almost entirely against spreading and influence benchmarks. We instead evaluate centrality against a *functional-diversity* reference—whether the genes a measure selects, as a set, cover the genome’s functional space or collapse it—a question the influence-benchmark tradition has not posed, and one that separates measures more sharply than any spreading benchmark.

We repurpose EDVS (Entropy of Degree-Vector Sums), a diversity measure introduced in scientometrics as an efficient, information-theoretic estimator of the variety, balance and disparity of a distribution [Zhao and Yang, 2023, Shannon, 1948, Cover and Thomas, 2006], as an annotation-free, topology-only centrality, and apply it to rice and *Arabidopsis* co-functional networks [Lee et al., 2015a, Obayashi et al., 2009, Lee et al., 2015b] and to yeast functional-association and physical protein-protein-interaction networks [Szklarczyk et al., 2023, Oughtred et al., 2021]. We establish three results and one bounded secondary one.

First, the criterion in routine use returns a functionally narrow candidate set, and an annotation-free audit makes this measurable. The audit is itself the instrument that reveals the deficit: we score a centrality against a *functional-diversity reference*—whether the genes it selects, as a set, cover the genome’s functional space or collapse it, relative to an annotation-count-matched maximum-entropy expectation—rather than against the spreading and influence benchmarks the field has used. The reference is reusable for any selection criterion. Audited this way, degree’s top-1% collapses functional coverage below the reference on every network tested: on the canonical rice network its selection spans an effective 15 pathways against the reference’s ~144, and 66 distinct pathways against EDVS’s 345. Three of our five networks are annotation-supervised by construction—the log-likelihood weights of RiceNet and AraNet were benchmarked against GO co-annotation, and STRING’s score draws on curated databases—so we report the audit stratified: the collapse is 1.6-3.3 bit there, and 0.5–1.1 bit on the two networks built without annotation (a co-expression and a physical-interaction network), with the same direction and the same ordering of measures. The deficit is amplified by that supervision but not created by it. Its explanation, however, is structural and follows from the rule: degree does not ignore bridging but returns the bridges that are also locally dominant—connector *hubs*—so a degree-ranked list holds a smaller share of the classical non-hub connector roles (18–19% on rice at the default resolution) than the genome itself does (26%), and it is that omitted class whose absence narrows the list’s functional span.

Second, EDVS recovers the omitted class. It isolates the high-participation, low-within-module-degree (*z*) pure connector that degree, Burt and betweenness all leave out: because its balance and disparity components penalise within-module concentration, its selections are structurally driven to low *z*, a mechanism we derive from the measure’s construction and confirm in every network and resolution tested—generalising across kingdoms (from the two plant networks to yeast) and across network genres (functional-association and physical protein-protein–interaction networks)—and it preserves the functional coverage degree loses.

Third, it does so without the community partition its own coordinates presuppose: the coverage that matters is carried by cross-module participation *P* —not, once degree is controlled, by *z*-and *P* cannot be computed without a partition; yet EDVS matches or exceeds top-*P* coverage on all five networks while using none, is invariant to the partition choice that moves *P*’s coverage by up to a bit across nine equally defensible alternatives, and retains 0.84 of its selection under an edge perturbation that leaves partition-based selections with 0.21–0.46. The axis so isolated is organizational rather than an importance signal: pre-registered probes—transcription-factor identity, essentiality, tissue-specificity, date/party-hub character, phenotype co-localisation—return null or reversed throughout, and a yeast test against a genome-complete essentiality label does not replicate even the plant pattern of importance tracking degree, so that coupling is itself network-dependent rather than fixed.

As a secondary result, the connector position carries a bounded, degree-independent, bidirectional cross-species footprint across the rice–*Arabidopsis* divergence. Underwriting the audit, we first quantify the degree bias of the curated knowledge that would otherwise be used to judge a degree-correlated detector—rendering such validation partly circular—which is why the reference we score against is annotation-matched rather than annotation-derived. The practical upshot is a concrete one for anyone ranking genes by centrality: the criterion in standard use returns a candidate set that is functionally narrow and structurally biased against the very cross-module genes such rankings are often meant to surface, and EDVS supplies an annotation-free, partition-free alternative that does not.

The class this leaves us with is defined by position rather than by importance. The higher-*z* connector hubs that betweenness returns remain locally dominant, whereas the low-*z* connectors EDVS isolates are broadly coordinating yet locally non-dominant. We characterise them here as a measurement result and no more: *why* genes come to occupy this position—its functional and evolutionary basis, and whether it maps onto any specific pathway or phenotypic role—is a question our results leave open rather than settle.

## 2 Results

We analysed the rice co-functional network RiceNet v2 and, to test whether the connector signature generalises beyond plant co-functional networks, two *Arabidopsis* networks (the independently constructed co-expression network ATTED-II and the same-framework AraNet v2) and two yeast (*Saccharomyces cerevisiae*) networks of deliberately different construction type: the functional-association network STRING (high-confidence, combined score ≥ 700, *N* = 5,716; and a medium-confidence sensitivity subset, ≥ 400, *N* = 6,290) and the physical protein-protein-interaction network BioGRID (*N* = 6,070). Throughout we compare EDVS against three reference centralities-degree, Burt’s structural-hole constraint, and betweenness— and, for analyses that contrast the genes *uniquely* selected by each measure (its “exclusive” set), we use the four-way definition throughout (each measure’s top-1% minus the top-1% sets of the other three; on the canonical rice network EDVS-exclusive *n* = 150, betweenness 82, degree 69, Burt 32), with a pre-registered top-2% sensitivity in yeast because its networks are smaller. Because the published RiceNet edge set spans a wide range of confidence (log-likelihood scores,LLS, 1–5) and its authors report higher precision for high-scoring links, we take as the canonical network the high-confidence subset (LLS ≥ 1.95; top 588,221 edges, ~33%; *N* = 20,851). All rice analyses are repeated on ~50% and full networks (Table 1).

**Table 1:** Core conclusions across the three rice confidence thresholds, grouped by the argument they serve. Qualitative results are invariant; quantities that shift with threshold are reported explicitly. Connector-share rows are at the default resolution (*r* = 1), where the role class is informative (Section 2.2); the *z*-gradient row is at the coarsest, where the four measures are furthest apart. Cross-kingdom (yeast) outcomes are summarised in Sections 2.4–2.6.

| Conclusion | Full (LLS>0) | ~50% | Canonical (~33%) |
| --- | --- | --- | --- |
| *The audit: what a degree-ranked list loses* |  |  |  |
| Degree collapses coverage below the reference (bits) | −1.60 | −2.44 | −3.26 |
| Degree distinct pathways vs EDVS | 186 vs 309 | 144 vs 301 | 66 vs 345 |
| Non-hub connector share: degree vs genome ($r=1$) | 19 vs 42% | 18 vs 29% | 18 vs 26% |
| Non-hub connector share: EDVS ($r=1$) | 78% | 63% | 55% |
| Burt→degree degeneration ($\rho$) | 0.995 | 0.994 | 0.993 |
| *Why curated knowledge cannot be the reference* |  |  |  |
| KEGG degree bias (member vs non-member median deg) | 60 vs 14 | 90 vs 17 | 59 vs 11 |
| Top-1% curated enrichment (Deg/Burt>EDVS) | yes | yes | yes |
| *The instrument: what EDVS recovers* |  |  |  |
| EDVS and betweenness not below reference | equiv. | equiv. | equiv.+ |
| Set-distinct pathways (EDVS≈BT>Burt>Deg) | yes | yes | yes |
| Within-module $z$ gradient (Deg>Burt>BT>EDVS, $r=0.5$) | yes | yes | yes |
| EDVS < betweenness on $z$ (3 resolutions) | 3/3 | 3/3 | 3/3 |
| EDVS lowest of the four arms (3 resolutions) | 3/3 | 2/3 | 2/3 |
| EDVS-exclusive inter-pathway edge fraction highest | yes | yes | yes |
| *Scope: the axis is organizational* |  |  |  |
| TF enrichment of EDVS-exclusive (deg-matched) | null (equiv.) | null (equiv.) | null (equiv.) |
| funRiceGenes recovery (EDVS, BT > deg-matched; Burt n.s.) | yes | yes | yes |

### 2.1 Why the audit cannot use curated knowledge: annotation is itself degree-biased

An audit needs a reference, but the obvious one—curated knowledge—is itself degree-biased, so scoring a degree-correlated criterion against it is partly circular. Genes in curated KEGG pathways are strongly enriched among high-degree nodes (Fig. 1A): rice pathway members (*n* = 4,137) have median degree 59 versus 11 for non-members (Mann–Whitney *p* ≈ 0, Cliff’s *δ* = 0.46). The most stringent degree/Burt selections are accordingly the most saturated with already-curated genes (top-1%: degree and Burt recover pathway members at 3.4× and 3.2× the genomic baseline—142 and 131 of 208—versus 2.9× for EDVS, 119 of 208); because EDVS requires no annotation, its selections escape that circularity. We treat curated-hit counts solely as evidence of annotation bias, not as a performance measure. This bias is not unique to plants: yeast shows it at comparable or larger magnitude (KEGG degree-bias Cliff’s *δ* = 0.39–0.59 across the three networks) with the same degree *>* Burt *>* EDVS gradient, so annotation-completeness does not by itself remove it.

**Figure 1.**
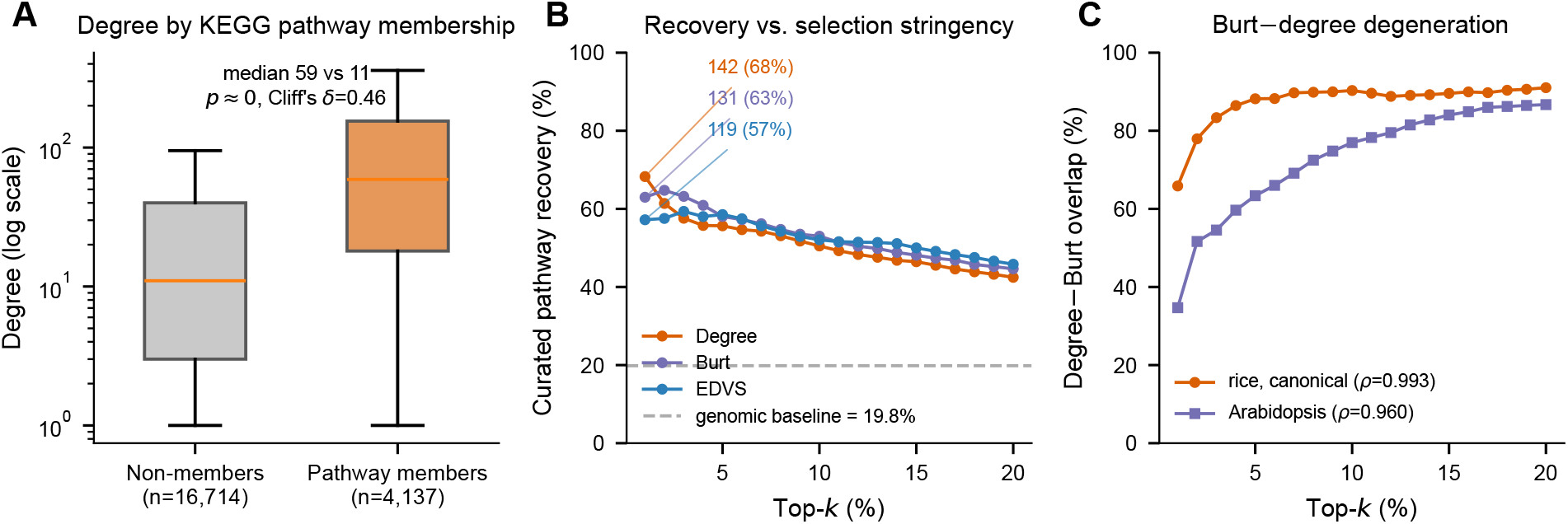
: Curated knowledge is degree-biased. (A) Degree of KEGG pathway members versus non-members (log scale). (B) Cumulative recovery of curated pathway members by degree, Burt and EDVS versus selection stringency, against the genomic baseline (dashed). (C) Top-*k* overlap of degree- and Burt-selected sets, rice versus *Arabidopsis*.

### 2.2 Audited against a functional-diversity reference, degree-ranked lists collapse and omit the non-hub connectors

We therefore score each selection not against curated knowledge but against an annotation-count-matched maximum-entropy reference—the rarefied Shannon entropy of a set’s propagated GO-BP annotations against a null matched on annotation count—asking a question the influence and spreading benchmarks do not pose: does the selected set, *as a set*, cover the genome’s functional space or collapse it? Audited this way, the criterion in routine use fails, and the four measures separate into two classes. Three of our five networks are *annotation-supervised* by construction: the log-likelihood edge weights of RiceNet v2 and AraNet v2 were benchmarked against GO-BP co-annotation as their gold standard, and STRING’s combined score includes database and text-mining channels drawn from curated pathways. On those networks the topology is not independent of the annotations we score against—modules resemble pathways partly because they were built to—so an audit run only there would push the circularity back a step rather than break it. Two networks are not annotation-supervised by construction: the *Arabidopsis* network used here is ATTED-II, whose edges are mutual-rank co-expression relations thresholded at *Z* ≥ 3.0, and BioGRID’s are physical interactions recorded by assay. No functional annotation enters either edge set, so on these two the reference really is external to the topology being scored. We therefore report the audit stratified, and treat those two as the independent replication that carries the claim. Degree- and Burt-selected sets fall far *below* this reference on every rice network (canonical effective pathways: degree 15, Burt 71, versus ~144, i.e. 2^7.17^ bits; canonical diff = −3.26 and −1.02 bits, both *p* ≈ 0), whereas EDVS and betweenness are never significantly below it (TOST-equivalent within ±0.15 bit or above). Consistent with a set-level rather than per-gene effect, EDVS-selected genes do not carry more GO terms each; rather their set covers far more distinct pathways (canonical: 345 EDVS, 339 betweenness, versus 220 Burt, 66 degree). The same ordering holds in *Arabidopsis* (degree and Burt below, diff = −0.54 and −0.19 bits; EDVS and betweenness equivalent).

This preservation extends cross-kingdom on functional-association networks but is bounded by network type. On yeast STRING the pattern reproduces strongly: degree-exclusive collapses hard (rarefied-entropy diff = −1.66 to −2.60 bits, larger than in rice) while EDVS- and betweenness-exclusive preserve; the small (*n* ≈ 5) top-1% Burt-exclusive set is unstable and its collapse is resolved by the pre-registered top-2% check. Set-distinct-pathway coverage keeps the connector-over-provincial ordering on every yeast network (EDVS 670/675/454 across the three networks versus degree 32/106/156).

The stratification separates the two halves of the argument. Ordered by construction, the degree collapse is −1.60, −2.44, −3.26 and −2.53 bits on the four annotation-supervised networks and −0.54 and ≈−1.1 bits on the two that use no annotation (ATTED-II and BioGRID)—roughly twice as large where the edges were themselves supervised by the annotations we score against, exactly as that construction would predict, but not an artefact of it, since the direction and ordering of the four measures are identical on the two networks built without annotation. The audit’s claim—that a degree-ranked list is functionally narrow—therefore holds on 5 of 5 networks and every construction type tested. The remedy’s claim does not travel as far: on the sparse BioGRID physical-PPI network the EDVS- and betweenness-exclusive sets themselves dip below the reference (≈ −0.25 and −0.35 bits), so while degree still collapses there by an order of magnitude more, EDVS no longer preserves coverage.

What explains the collapse is visible in the composition of the selections themselves. We describe each top-1% list by the share of its members occupying the classical non-hub connector roles (R3/R4 of the Guimerà-Amaral cartography: *z <* 2.5, *P >* 0.625), alongside the share of the genome in those roles. This is a description of what each criterion returns, not an enrichment test against a null-degree cannot meaningfully be degree-matched against itself—so we report proportions and read them as composition. At the default resolution on the canonical rice network, where 26% of the genome occupies those roles, a degree-ranked list holds 18%; on the five networks the shares are 18, 18, 19, 2 and 0% against genome shares of 26, 29, 42, 21 and 7%. EDVS’s lists hold 55, 63, 78, 64 and 44%, Burt’s 30, 38, 43, 7 and 4%, betweenness’s 38, 45, 55, 22 and 23%. This follows directly from the selection rule: at this resolution 28–36% of degree’s rice selections do clear the participation cutoff, yet only 18–19% qualify as R3/R4, because the role additionally requires *z <* 2.5 and degree’s picks are module hubs. Degree is not blind to bridging—it returns the bridges that are also locally dominant, connector *hubs* (R6/R7)—and leaves the non-hub connectors out. The collapse in Fig. 2 is the measurable consequence; this composition is the reason for it, and the omitted class is the one the next section shows EDVS recovers.

**Figure 2.**
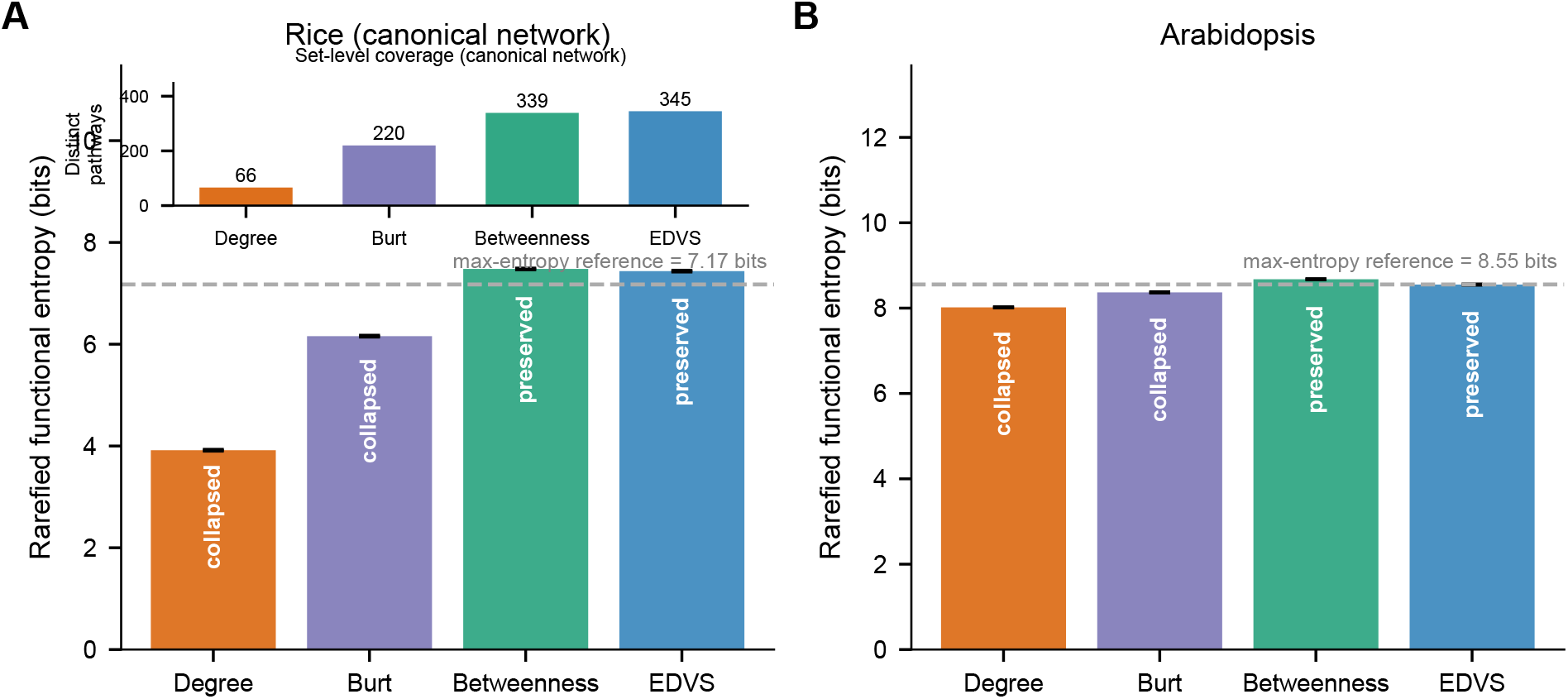
: A degree-ranked list is functionally narrow; connector-type selections are not. Rarefied GO-BP functional entropy of each of the four selected sets against the annotation-count-matched maximum-entropy reference (dashed), for rice (canonical) and *Arabidopsis*. Degree’s selection falls furthest below the reference and Burt’s follows it; EDVS and betweenness sit at or above it. The deficit is stratified by construction and holds on the networks built without annotation (Section 2.2). Inset: set-level distinct pathways (EDVS 345, betweenness 339, Burt 220, degree 66, canonical). Error bars: bootstrap 90% CI.

Two resolution-related scope statements apply here. The participation coordinate has no absolute meaning independent of the number of modules: at the coarsest resolution tested the connector roles comprise 0.004–4.4% of the genome and *no* measure’s selection clears the classical cutoff (degree 0%, EDVS 0.7–2% on the plant networks), while at the finest they expand to 11–63% and the label largely stops discriminating—at which point degree’s rice selections do reach baseline (1.2– 1.7×). The depletion we report is therefore stated at the coarse and default resolutions, where the class is informative, and we report all three (Fig. 3A shows both the default and the coarsest resolution for this reason). Second, this analysis scores agreement with a role definition that itself requires a partition; we use it as an external reference for what each criterion selects, not as a ground truth about the network.

**Figure 3.**
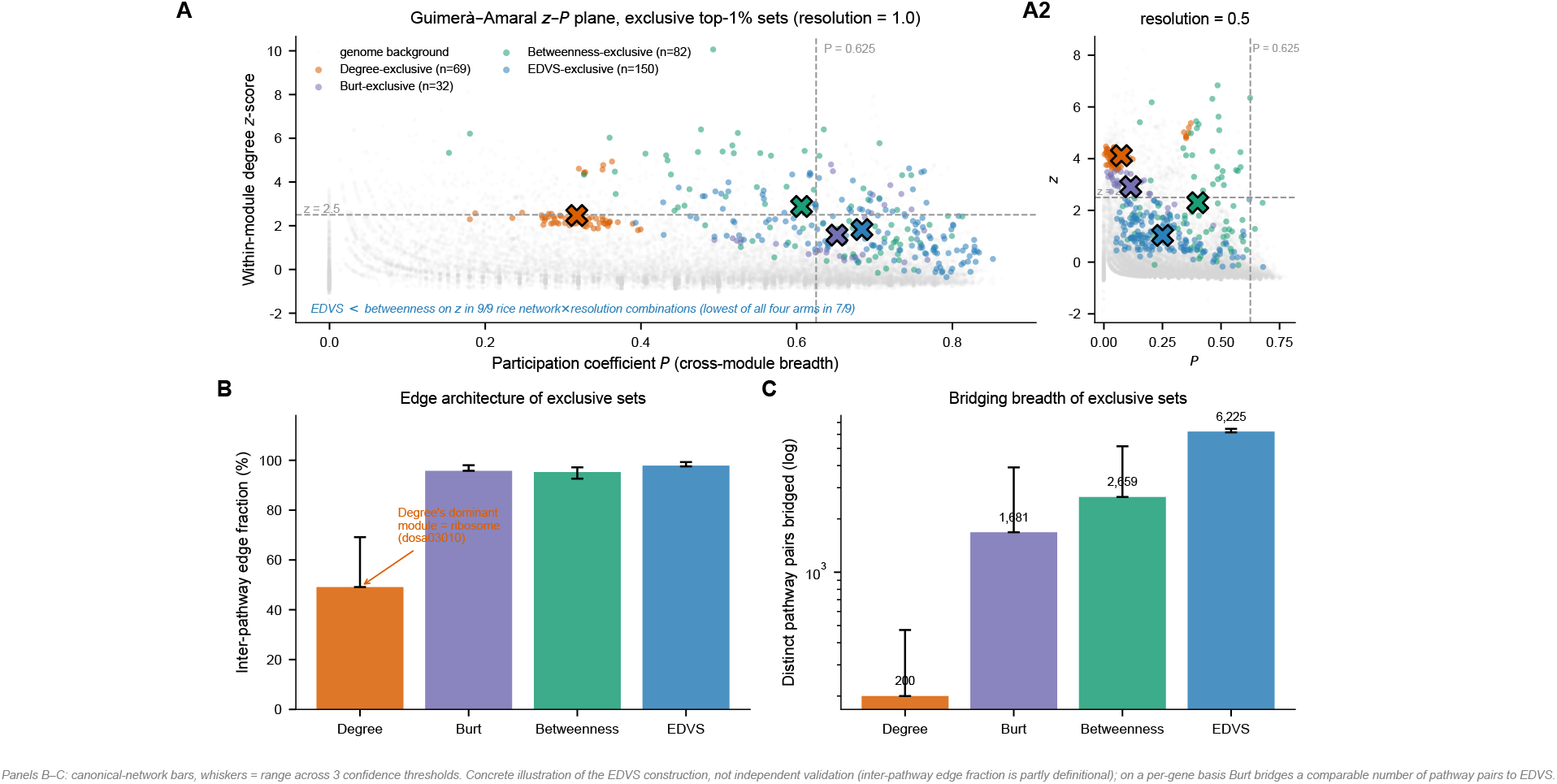
EDVS recovers the connector class a degree-ranked list omits. (A) Guimerà –Amaral within-module degree (*z*) versus participation coefficient (*P*) for the four exclusive top-1% sets at the default resolution (*r* = 1), with the coarsest resolution (*r* = 0.5) alongside: the participation cutoff is not an absolute scale, and at *r* = 0.5 no measure’s selection clears it, degree included. The measures form a monotone *z*-gradient degree *>* Burt *>* betweenness *>* EDVS; EDVS occupies the low-*z ex*treme, with betweenness’s selections sitting systematically above it on *z* (9/9 network × resolution combinations). (B) Inter-pathway edge fraction per exclusive set. (C) Distinct pathway pairs bridged per set. Four-way exclusive sets; canonical rice EDVS-exclusive *n* = 150, betweenness 82, degree 69, Burt 32.

### 2.3 EDVS recovers the omitted class: the lowest-within-module-degree connector, distinct from betweenness

What degree omits, EDVS recovers. Among the four classical measures compared here, EDVS alone isolates the region of the centrality space that the audit implicates. Placing all four measures on the Guimerà –Amaral within-module degree (*z*) / participation coefficient (*P*) plane, they form a monotone gradient in within-module *z*–degree 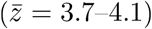 *>* Burt (2.6–2.9) *>* betweenness (2.3–2.4) *>* EDVS (0.9–1.1) at the coarsest resolution—with EDVS at the pure-connector extreme. EDVS has the lowest within-module *z* of all four arms in seven of nine rice network × resolution combinations; the two exceptions are the small (*n* = 32-39) Burt-exclusive set at the intermediate resolution, which dips marginally below it (mean *z* 1.64 vs 1.97 and 1.43 vs 1.48)—the same small-set instability we disclose in yeast (Section 2.4), and one that does not place Burt in the connector quadrant, since its selections remain at low participation. The contrast that is both robust and fully consistent is with betweenness, the measure that competes for the connector role: EDVS’s selections sit at lower *z* than betweenness’s in all nine combinations, significantly throughout (9/9; *p* = 1.3 × 10^−8^ to 1.1 × 10^−18^), so betweenness’s selections sit systematically above EDVS’s on *z*. The *P* axis separates them in the same direction but less uniformly: EDVS’s participation exceeds betweenness’s in 12 of 15 network × resolution combinations, the three exceptions all falling at the coarsest resolution and on only three of the five networks. The distinction is therefore that **EDVS isolates a purer connector class than betweenness:** where betweenness returns connector-*hubs* that bridge modules yet remain locally central, EDVS returns broadly bridging connector *nodes* that dominate no single module.

This connector identity is visible in the genes’ edges—partly definitional, an illustration of what the measure selects for rather than independent validation of it. On the canonical network almost every edge of an EDVS-exclusive gene bridges two different KEGG pathways (97.9%, the highest of the four arms; 97.6-99.3% across networks), whereas only 49.1% of a degree-exclusive gene’s edges are inter-pathway—the remainder concentrating within a single dominant module that is, repeatedly, the ribosome (dosa03010), a core essential process. In aggregate, EDVS-exclusive genes bridge far more distinct pathway pairs (6,225) than betweenness (2,659), Burt (1,681) or degree (200). EDVS-selected genes remain moderately high-degree (median degree at the 95.5th percentile genome-wide; *p <* 10^−100^) and high-betweenness, so they are genuine topological bridges, but they are markedly less degree-extreme than degree/Burt selections (median degree ~ 2.5× lower) and overlap the degree top-1% by only 2–5%. Although EDVS is rank-correlated with degree genome-wide (Spearman *ρ* = 0.88)-expected, since degree bounds the richness component of any diversity measure and is a *constituent* of EDVS rather than a competitor—the operative top-1% selections diverge sharply, and EDVS’s distinctive signal is the degree-independent low-*z* increment. It recovers this connector signal without the all-pairs shortest paths of betweenness (*O*(*nE*)) or the community detection that participation requires; capturing a related but distinct property, it is correlated with but not redundant to betweenness (Spearman *ρ* = 0.57–0.72).

A sharper version of the same question applies to the entropy itself. Because EDVS(*i*) = *H*(*M* [*i*, :]) and the support of *M* [*i*, :] is the two-hop neighbourhood *N*_2_(*i*), EDVS is bounded by log |*N*_2_(*i*)|, raising a natural null: that the measure is merely two-hop reach—*variety* alone, without the balance and disparity terms Mechanism 1 invokes. Genome-wide this looks serious (log |*N*_2_| and EDVS: Spearman *ρ* = 0.92-0.98 across the five networks), but the operative selections diverge completely, in the direction the mechanism predicts. The two top-1% sets barely intersect (Jaccard 0.026–0.118, never above 0.12) and occupy opposite regions of the *z*–*P* plane: at the coarsest resolution the |*N*_2_|-selected set sits at median *z* = 4.22, *P* = 0.11 with 90% above the hub cutoff, versus EDVS’s *z* = 1.16, *P* = 0.24 with 6.7% above it (canonical rice; the same contrast holds on all five networks). A variety-only criterion thus selects *provincial hubs*, not connectors-it is the balance and disparity terms, not support size, that move the selection cross-module. EDVS is significantly lower-*z* than the |N_2_| baseline in 13 of 15 network × resolution combinations; the two exceptions (yeast, *k* = 57, at the two finer resolutions) shrink toward but miss significance (*δ* = −0.18, −0.08).

This pattern—high genome-wide correlation with a barely-intersecting tail—recurs for every reference measure (degree *ρ* = 0.88 with 2–5% top-1% overlap; log |N_2_| *ρ* = 0.92–0.98 with Jaccard ≤ 0.12), inviting the objection that such a tail is noise rather than signal. Two observations argue otherwise. First, EDVS’s selection is itself robust to perturbation at exactly this scale: under 10% random edge removal its top-1% set retains a Jaccard of 0.84 against the unperturbed selection on both real networks (0.843 yeast, 0.837 rice; *n* = 10 and *n* = 5 replicates), against 0.21–0.46 for participation-based selections under the identical procedure-a tail keeping five si ths of its membership under perturbation is not rank jitter. Second, the disagreement is systematic, not random: the two tails sit in different, reproducible regions of the *z*–*P* plane rather than scattering, which shared-ranking noise cannot produce. The |N_2_| baseline is best read in this light: its top-1% selections overlap degree’s substantially (Jaccard 0.36–0.58 on rice and yeast), 5–100× more than EDVS’s (0.00–0.04)—a variety-only criterion largely reselects what degree selects, the same degeneration Burt’s constraint shows at the selection level (Jaccard with degree 0.45–0.70 on those networks, *Arabidopsis* the shared exception at 0.09 and 0.21). We therefore read |N_2_| not as an independent variety baseline but as a demonstration that variety without balance and disparity collapses back toward degree.

Placing the same selections in the role cartography that originally defined the connector concept [Guimerà and Amaral, 2005] corroborates this externally. Under its thresholds, EDVS’s selections are the most concentrated of the four measures in the non-hub connector roles (R3/R4: *z <* 2.5, *P >* 0.625) at every resolution where those roles remain informative: at the default resolution 44– 78% of an EDVS list occupies them, against 0–19% of a degree list and genome shares of 7–42% (Section 2.2). We report these shares rather than recall, which is uninformative here by construction: the role class contains thousands of genes while a top-1% selection contains a few hundred, capping recall near a few per cent for every measure regardless of quality-which is also why these shares do not conflict with the low F1 scores of partition-free detectors on synthetic benchmarks (Supplementary Material): a planted connector set small enough for recall to be meaningful is not the regime a thousands-of-genes role class presents, where precision is the only informative half of the pair. EDVS does this without the partition the role definition itself requires: role labels are strongly partition-dependent, with only 26.5% of nodes retaining the same role across the three resolutions on the largest network. EDVS therefore *recovers* the classical non-hub connector role partition-free—recovery of the role, not an identity between measure and role.

### 2.4 The low-*z* connector generalises across kingdoms and network genres

To ask whether this is a plant / co-functional-network phenomenon or a general property of the measure, we repeated the *z*–*P* analysis in yeast on two deliberately different network types (Fig. 4). The result is the strongest cross-kingdom signal in this study: EDVS’s separation from betweenness on within-module *z* holds in *every* yeast network × set-size combination (6/6; one-sided ann-Whitney, |Cliff’s *δ*| = 0.27 to 1.00, *p* from 3.1 × 10^−2^ down to 3.9 × 10^−14^), and EDVS is the lowest-*z* of the four arms in 5/6 (failing only at the smallest set size on high-confidence STRING, where an *n* = 5 Burt-exclusive set dips marginally below EDVS at two of three resolutions, resolved by the pre-registered top-2% check). Crucially this holds on the physical PPI network as well as the functional-association one–on BioGRID, EDVS’s mean within-module *z* is actually negative (−0.15 to −0.33), an even purer connector than in the plant networks. EDVS’s structural signature-the lowest-*z* cross-module connector, sharply distinct from the higher-*z* betweenness arm—therefore generalises from rice to yeast and from functional—association to physical-interaction networks.

**Figure 4.**
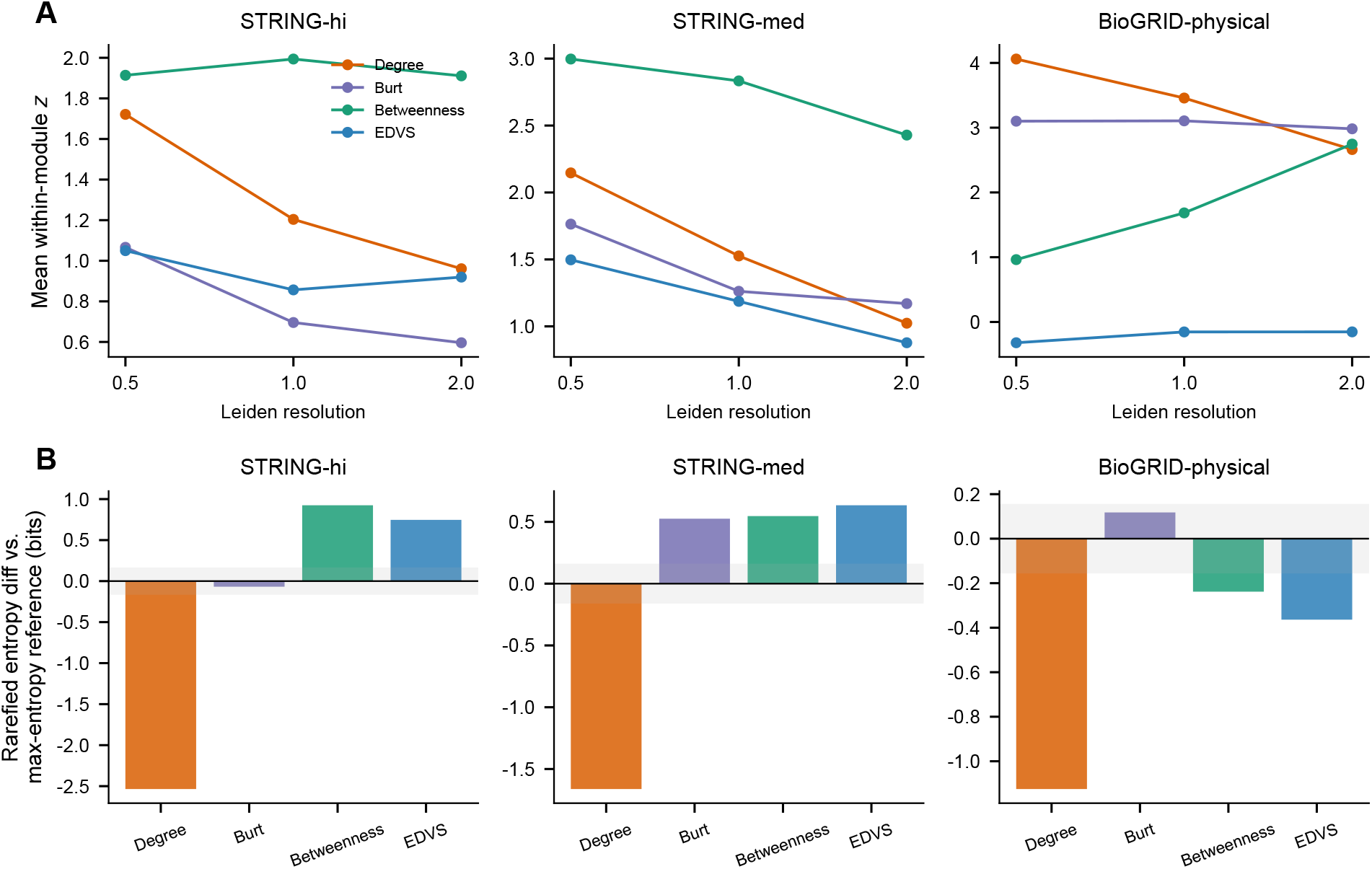
The low-*z* connector generalises to yeast across network genres. (A) Four-arm within-module-*z* gradient across the three yeast networks (STRING high-confidence, STRING medium-confidence, BioGRID physical PPI); EDVS is lowest-*z* and its separation from betweenness holds in 6/6 network × set-size combinations. (B) Functional diversity preserved by EDVS and betweenness on STRING and attenuated on the sparse BioGRID physical-PPI network.

### 2.5 The coverage is carried by participation, and EDVS attains it without a partition

The audit of Section 2.2 is a two-class result; we next asked what, on a continuous scale, drives the coverage it measures, by sliding a selection window along each purity axis at fixed set size (Fig. 5A). Read naively, coverage *rises* with within-module degree *z* in all 15 network × resolution combinations-the opposite of the expected direction—but this is a degree artefact: mean degree rises steeply along the same *z* windows (~1 to ~75), and coverage rises with degree for trivial reasons. Once degree is controlled the picture inverts. In a partial-effects regression of coverage on (purity, degree), participation *P* retains a significant positive effect in 11 of 15 combinations, while the apparent *z* effect vanishes or reverses in the plant networks; a non-parametric matched-degree-bin check agrees in 14 of 15. By combination, 10 of 15 show the clean pattern (*P* drives coverage, *z* is a degree shadow), 4 of 15 show no significant effect for either a is once degree is removed (both rice_full_ resolutions and two yeast conditions), and 1 of 15—yeast at the finest resolution-shows an independent, stronger *z* effect, reported rather than excluded. The axis carrying functional representativeness is therefore cross-module participation, not local dominance. One boundary is explicit: while *P drives* coverage, the pre-registered mechanism for *how*-that higher *P* raises coverage by spanning more functional modules—was not confirmed (1 of 15 combinations met all mediation criteria, in a suppression rather than mediation pattern), so we claim the driver, not the route.

**Figure 5.**
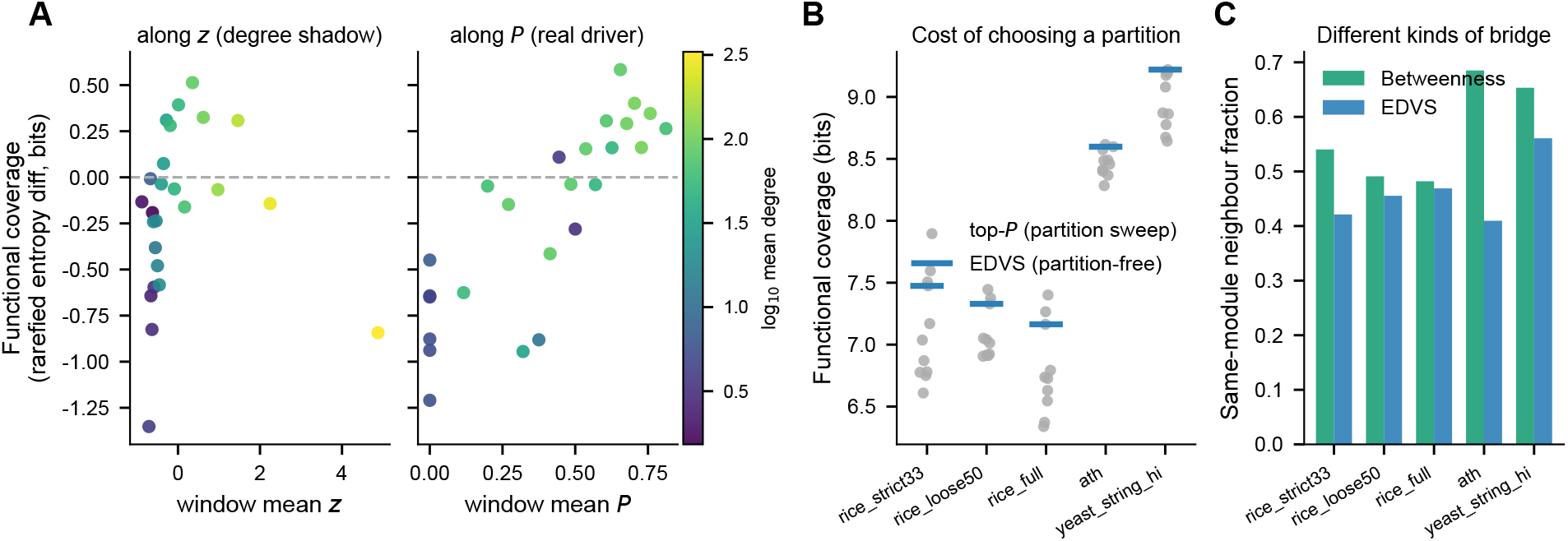
Functional coverage is driven by participation, and EDVS attains it without a partition. (A) Functional coverage of a fixed-size selection as the selection window slides along within-module degree *z* (left) and along participation *P* (right); points coloured by the window’s mean degree. Coverage appears to rise with *z*, but mean degree rises with it—the effect is a degree shadow and does not survive degree control, whereas *P* ‘s effect does (rice canonical network, resolution 1.0; 11/15 network × resolution combinations for *P*). (B) Functional coverage of the top-*P* selection under nine reasonable partitions (Leiden/Louvain at three resolutions, Infomap under three seeds) versus EDVS, which uses no partition and is invariant: *P* ‘s coverage swings by 0.33-1.06 bit. (C) Fraction of a selected gene’s neighbours lying inside its own module: betweenness-exclusive genes keep more neighbours at home, EDVS-exclusive genes bridge outward.

One could ask whether the same coverage is reachable by selecting directly on the low-*z*, high-*P* connector coordinates rather than through EDVS—but only after first computing *z* and *P*, which requires the very partition EDVS avoids. The comparison that follows therefore asks not whether EDVS reaches a region a partition-based rule cannot, but whether it reaches the same region without one. We pre-registered the expectation that *P*, being selected directly on cross-module participation, would win. It did not. At matched set size, EDVS matched or exceeded the coverage of a top-*P* selection on all five networks (coverage ratio 1.01–1.06; set-level distinct pathways 1.19–1.3 ×): EDVS is not a degraded approximation of *P*, but reaches *P*-level coverage without a partition. The cost of that partition is then directly visible (Fig. 5B). Recomputing the top–*P* selection under nine reasonable partitions—Leiden and Louvain at three resolutions, Infomap under three seeds-*P* ‘s coverage swings by 0.33–1.0 bit on every network, while EDVS, using no partition, is invariant by construction. We tested a further prediction, that EDVS would also retain coverage better under edge noise, and report it as unsupported outside yeast: under 10–30% edge addition and deletion, EDVS and *P* retained coverage indistinguishably on the rice and *Arabidopsis* networks (0.97–1.05 for both), and only on yeast was *P* measurably more fragile (mean retention 0.906 versus 1.006). Because this round used a single perturbation per noise level, we report the plant result as *not resolved* rather than as evidence of equivalence.

Finally we asked whether the lower-*z* purity separating EDVS from betweenness (Section 2.3) is a distinction with structural consequences or merely a coordinate difference, contrasting the genes each measure uniquely selects with all directions fixed in advance. Two of three pre-registered signatures hold. EDVS-exclusive genes have significantly lower *z* and higher *P* than betweenness-exclusive genes on all five networks (most *p <* 0.001) and—more concretely—a significantly smaller fraction of their neighbours inside their own module on all five networks (Fig. 5C), with neighbours spread across more distinct functions on four of five. Betweenness returns a bridge that brings its own module along; EDVS returns a bridge whose neighbourhood sits elsewhere. The third signature does not hold: at matched pathway coverage we found no support, at the primary threshold on any network, for the expectation that the betweenness-selected set is functionally more skewed and the EDVS-selected set more even (a partial recovery at a 2% threshold on the rice networks is a threshold-edge effect and we do not treat it as support); one neighbour-dispersion operationalisation also failed and is reported rather than dropped. The two measures therefore select different *kinds* of connector—differing in node-level topological purity and neighbourhood composition, but not in the functional-distribution shape of the selected set. We report a difference in kind, not in quality.

### 2.6 The connector axis is organizational, not an importance signal

The genes EDVS selects are organizationally central but are not, in the plant networks, the network’s *important* genes in the essentiality sense. Under the four-way definition, EDVS-exclusive genes (*n* = 150) are not enriched for transcription factors (TF proportion 1.33%, at or below the ~5. % genomic baseline; degree-matched *z* = −0.99, and *z* = −0.63 to −1.17 across rice networks, all n.s.), while degree- and Burt-exclusive sets contain no TFs at all (0 of 69 and 0 of 32). In *Arabidopsis*, degree-exclusive genes are enriched for embryo-essential (*EMB*) genes (4.69%; OR= 5.03, *p* = 0.0018) whereas EDVS-exclusive genes are not (1.60%; OR= 1.63, *p* = 0.44). One external label does separate connector-type from provincial selections: EDVS-exclusive genes are enriched, beyond a degree-matched null, for genes already functionally characterised in rice (funRiceGenes: 36.7% on the canonical network, *z* = 3.99; 31.8–36.7%, significant on all three networks), an effect shared with betweenness (39.2–48.8%, significant on all three) but not with Burt (n.s. on all three, *z* = −0.44*/*1.15*/*1.21) or degree (below the null). Because funRiceGenes catalogues already-studied genes, this reflects research attention—connector-type selections reach more of the already-characterised genome than degree does—rather than importance.

Yeast, whose genome-complete essential-ORF set removes the label sparsity that limits the plant tests, qualifies this dissociation sharply: the essentiality contrast *does not replicate* on either S RING network (Fig. 6C). We report the pre-registered top-2% sets here, rather than top-1%, because the yeast networks are small and top-1% exclusive sets are correspondingly unstable—a sensitivity we pre-registered for exactly this reason. Under a degree-matched null, degree-exclusive genes are essentiality-*depleted* rather than enriched (high-confidence STRING: 13.6% of *n* = 22 versus a 38.9% expectation, *z* = −2.48; medium-confidence: 28.9% of *n* = 38 versus 42.3%, *z* = −1.76), while EDVS-exclusive genes sit at the null (38.3% of *n* = 60 versus 34.5%, *p* = 0.31; 30.4% of *n* = 79 versus 27.8%, *p* = 0.35). The plant pattern—importance tracking the degree a is—therefore does not hold here; we describe this as a failure to replicate rather than as a reversal, since EDVS-exclusive genes are not significantly essentiality-enriched in either network. The corresponding top-1% figures, where the degree-exclusive set falls to *n* = 10, are given in the Supplementary Material and are not the basis of any claim. A complete label is not the same as a powerful test: while yeast removes the label sparsity that limits the plant essentiality analysis, the exclusive sets themselves remain small (*n* = 22–121), and the equivalence assessment in Section 2.7 accordingly finds these particular probes inconclusive rather than establishing an absence. This contrast was pre-registered as a two-arm test (degree-versus EDVS-exclusive); Burt- and betweenness-exclusive sets were not part of it, and the exploratory figures for those arms are reported in the Supplementary Material and flagged as such. On the physical-PPI network the raw degree—lethality signal does not survive degree matching. Taken with the plant results, this shows the coupling between centrality and importance is itself *network-dependent* rather than fixed—a caution against reading importance directly off any single centrality, consistent with this paper’s opening premise that centrality conflates importance with organizational role. Whether the organizational axis EDVS isolates carries importance is therefore a property of the network, not of the measure, and in the yeast test it does not.

**Figure 6.**
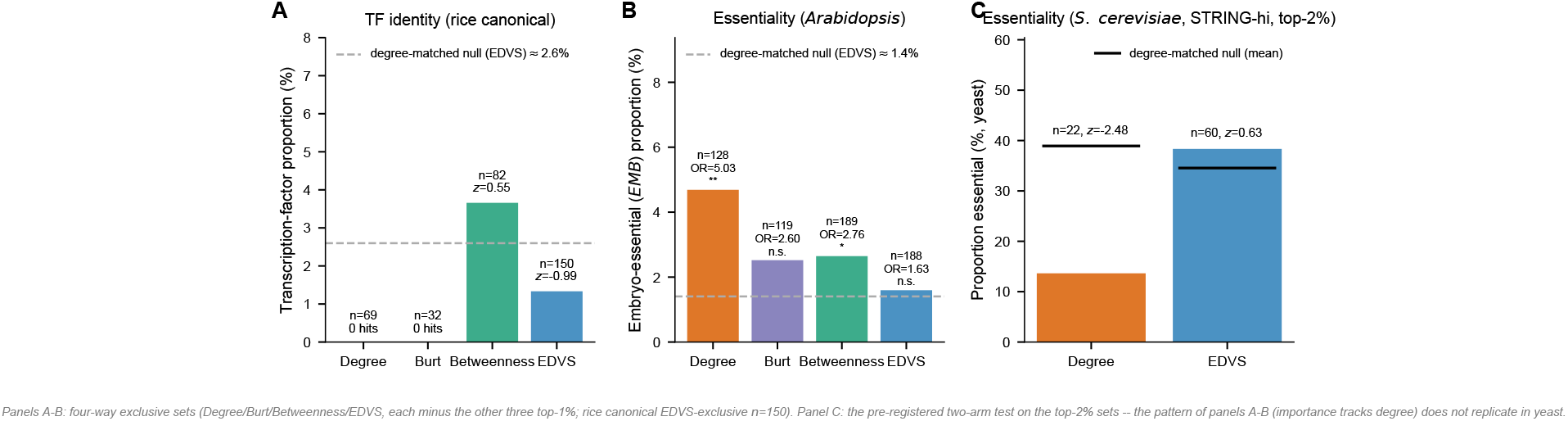
The connector axis is organizational, not an importance signal. (A) Transcription-factor proportion of each four-way exclusive set against the genomic baseline with a degree-matched null; EDVS-exclusive genes are not TF-enriched. (B) Embryo-essential (*EMB*) enrichment in *Arabidopsis*: elevated for degree-exclusive, not EDVS-exclusive. (C) Yeast essentiality (genome-complete set, high-confidence STRING, pre-registered top-2% sets): under a degree-matched null the plant pattern does not replicate—degree-exclusive genes are essentiality-depleted (13.6% of *n* = 22 vs a 38.9% null, *z* = −2.48) and EDVS-exclusive genes sit at the null (38.3% of *n* = 60 vs 34.5%, *p* = 0.31) —evidence that the centrality—importance coupling is network-dependent. Panel C is the pre-registered two-arm test; panels A-B use four-way exclusive sets (canonical rice *n* = 150).

### 2.7 The axis is organizational: where testable, its independence from importance is an equivalence, not an absence of evidence

Consistent with the axis being organizational rather than an importance signal, a set of pre-registered probes for a phenotypic or importance signal on the EDVS connector class returned null or reversed results throughout, each against a degree-matched null: yeast and *Arabidopsis* essentiality (Section 2.6); transcription-factor identity; tissue-specificity of expression (EDVS-exclusive genes are, if anything, more tissue-specific, not less); date-hub versus party-hub character (EDVS connectors sit at the genome baseline of neighbour co-expression—markedly less “party” than provincial hubs, but not affirmatively “date” against random); co-localisation with rice yield/stress Q s; and co-association across rice yield and abiotic-stress QTLs as a trade-off proxy (no enrichment; if anything the provincial arms, not EDVS, trend higher).

A set of non-significant tests does not establish an absence, so we asked which of these outcomes are demonstrated absences and which are merely undetected, applying to each probe the same two one-sided equivalence procedure we use for functional coverage (margin ±5 percentage points, 90% intervals). The probes separate into three kinds. For transcription-factor identity on all three rice networks, and for embryo-essentiality in *Arabidopsis*, EDVS-exclusive sets are statistically *equivalent* to their degree-matched nulls (differences −1.25 to +0.23 pp; intervals contained within the margin, of tests): under the post-hoc margin defined in Section 4.9, these are demonstrated absences, not failures to detect, and they are what licenses the claim that the axis is separable from importance rather than merely uninformative about it. For yeast essentiality the equivalence test is *inconclusive* rather than supportive: with *n* = 36-121 the intervals span roughly ±10 pp and straddle both the null and the margin, so we report those probes as underpowered and draw no absence claim from them. One probe is not null at all: EDVS-exclusive sets are genuinely enriched for already characterised rice genes (funRiceGenes; +8.5 to +13.5 pp, intervals excluding both zero and the margin), a precisely estimated excess that inde es research attention rather than importance (Section 2.6).

We therefore report a bounded scope statement: the class EDVS isolates is defined by organizational position; for two importance markers we can state its independence as an equivalence, for the remainder the current public references leave the question underpowered, and in no case do we find it to be a marker of phenotypic importance. Full pre-registered protocols and per-probe statistics are in the Supplementary Material.

### 2.8 The connector position carries a bounded, degree-independent cross-species footprint

As a secondary result, the connector position leaves a degree-independent evolutionary footprint across the rice—*Arabidopsis* divergence. Using RiceNet v2 and the same-framework AraNet v2 with 1:1 orthologues (Fig. 7), the *Arabidopsis* orthologues of rice EDVS-exclusive connectors sit at elevated AraNet EDVS percentiles relative to a rice-degree-matched null (EDVS shift = +6.31 points, *z* = 4.97, *p* ≤ 0.0005; bidirectional, ath→rice = +14.44, *z* = 9.23), while the paired degree-percentile shift is negligible (+0.62, *p* = 0.34), so the role-specificity contrast excludes zero (Δ = +5.69, 90% CI [4.76, 6.65]). Because AraNet and RiceNet share a construction framework we treat this as an *upper* bound (Cliff’s *d* ≈ 0.82); the independently constructed ATTED-II gives a method-independent *lower* bound where the signal persists but attenuates (*z* = 2.16, *p* = 0.011, *d* ≈ 0.20). We therefore report the position as boundedly and bidirectionally conserved, not strongly conserved. A sequence-level check (percent protein identity, 69. % versus 63.3% degree-matched background, *p* = 0.002, *n* = 52; dN/dS unavailable at the monocot–eudicot distance) is corroborating rather than decisive.

**Figure 7.**
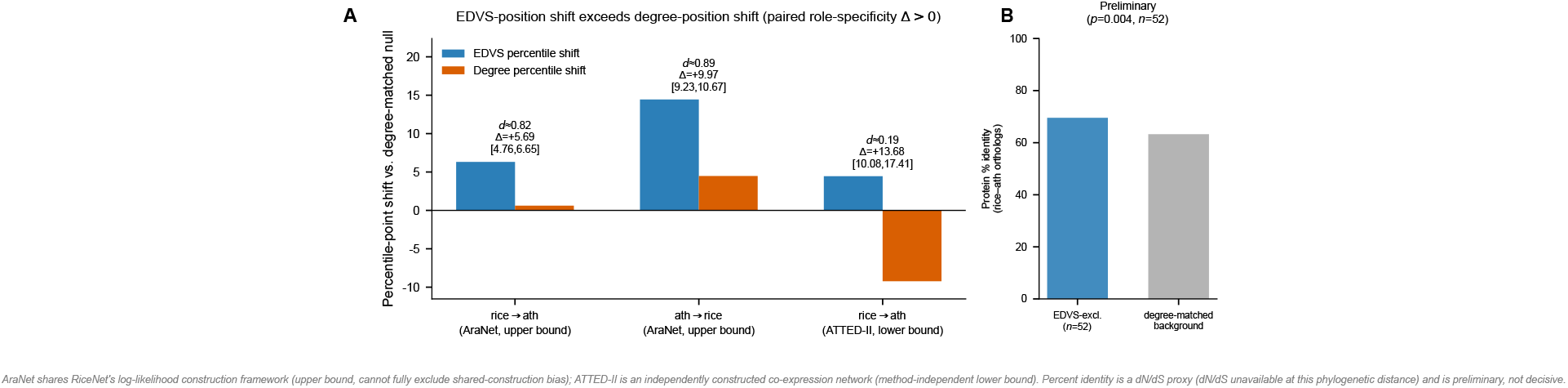
Connector position is boundedly, degree-independently conserved across the rice—*Arabidopsis* divergence. EDVS-percentile shift of the orthologues of rice EDVS-exclusive connectors relative to a degree-matched null, in the same-framework AraNet v2 (upper bound) and the independent ATTED-II (lower bound), with the paired degree-percentile shift for contrast; both directions shown. Focal set *n* = 125; preliminary protein-identity panel (B) uses *n* = 52.

### 2.9 Robustness across confidence thresholds and networks

Both halves of the audit hold across all three rice confidence thresholds: a degree-ranked list collapses functional coverage relative to the reference and holds a smaller share of non-hub connectors than the genome does, while EDVS preserves the coverage and recovers the omitted class—the latter result also across kingdoms and network genres. The importance/organization dissociation holds in the plant networks but, as Section 2.6 reports, does not replicate on the yeast essentiality test, which we disclose rather than select around.

## 3 Discussion

Our results recast the search for coordinating genes as a measurement problem. Centrality is used to rank genes, yet what a ranking omits is rarely asked. Scored against a functional-diversity reference, the criterion in standard use returns a narrow list: degree’s candidate set collapses functional coverage by up to 3.3 bit, on all five networks and on every construction type tested—including the two whose edges owe nothing to functional annotation, where the effect is smaller but of the same sign and ordering. An EDVS-ranked list does not collapse, and the reason it does not is the substance of this paper: EDVS recovers a class of genes that degree systematically leaves out. Named in the established vocabulary of the Guimerá—Amaral cartography, these are the non-hub connectors—low within-module degree (*z*), high cross-module participation (*P*): genes that bridge modules without dominating any one of them. Degree misses them not because it ignores bridging but because it returns the bridges that are *also* locally dominant—connector hubs. The composition of the lists makes the omission concrete: where the genome holds 26% of its genes in those roles, a degree-ranked list holds 18% and an EDVS-ranked list 55%. Importance and organizational role are separable properties, and degree, by construction, resolves the tension between them in favour of importance.

The asymmetry between the two halves of this result matters. The deficit is general: every network, every construction, every threshold. The remedy is bounded: EDVS preserves coverage on functional-association and co-expression networks but not on the sparse physical-interaction network, where degree still collapses by an order of magnitude more but EDVS no longer holds the reference. A blind spot that is everywhere and an instrument that works in most places is the honest shape of the finding.

That EDVS occupies the low-*z* connector region follows directly from its construction. Because EDVS is the Shannon entropy of a node’s aggregated neighbourhood connectivity, its balance component penalises connectivity concentrated within a single module and its disparity component penalises the redundant, mutually similar neighbourhoods that within-module concentration produces; both drive its selections toward cross-module spread and therefore toward low *z*. Degree counts edges without regard to their modular distribution, and betweenness rewards shortest-path centrality, which in dense modular networks accrues to locally dominant nodes; EDVS alone takes as its objective a quantity that is maximised by cross-module spread and minimised by local dominance. The construction fixes the direction of this effect, and the empirical within-module-*z* gradient— EDVS below betweenness in every network and resolution tested, and lowest of the four measures in most of them (the small Burt-exclusive set dips marginally lower at one intermediate resolution), now confirmed across kingdoms and network genres including yeast—fixes its magnitude.

Two of the contributions are, to our knowledge, new. The first is the isolation of the low-*z*, high-participation connector as a named, recoverable class, obtained without annotation and without community detection and stable from rice to yeast across both functional-association and physical-interaction networks. The connector-versus-provincial distinction is itself an older result [Guimerá and Amaral, 2005]; what EDVS adds is an annotation-free, partition-free way to isolate the connector’s low-*z* extreme—a region degree and constraint leave out as provincial and that betweenness leaves out as a higher-*z* connector hub. The *z*-P cartography is descriptive vocabulary, not a competitor EDVS is scored against: it names and locates the class EDVS reaches by local neighbourhood aggregation alone. The second contribution is the evaluation paradigm—assessing a centrality against a functional-diversity reference—which the influence- and spreading—benchmark tradition had not applied, and which separates connector-type from provincial-hub measures more cleanly than any spreading benchmark. Preserving functional diversity is a property EDVS shares with betweenness and marks the connector class as a whole; EDVS’s distinctive signature is the *z*-axis. Burt’s constraint sits between the poles—connector-leaning in its structural-hole construction [Burt, 1992, 2004], provincial in its diversity collapse and, as we find empirically, in its degeneration onto degree—and we report that gradient rather than forcing a binary.

The class EDVS isolates is organizational, defined by position rather than by importance. In the plant networks the provincial hubs degree selects concentrate within single dominant modules, repeatedly the ribosome, and are enriched for essential genes and transcription factors; the connectors EDVS selects are neither, so that importance in the essentiality sense there tracks the degree a is. Yeast, whose genome-complete essential-gene set removes the label sparsity of the plant tests, qualifies this sharply. The classical centrality—lethality relation [Jeong et al., 2001] does not replicate on either functional-association network: degree-exclusive genes are essentiality-depleted rather than enriched, and EDVS-exclusive genes sit at the degree-matched null. The coupling between centrality and importance is therefore network-dependent rather than fi ed. The one external label the connectors recover more than degree—already-characterised rice genes—is shared with betweenness but not with the provincial arms, and indexes research attention rather than importance. We therefore do not present the connector class as a marker of importance.

The connector position does, however, leave a degree-independent evolutionary footprint. Across the rice—*Arabidopsis* divergence the position is conserved, degree-independently and in both directions. Its magnitude is method-dependent, bracketed between a same-framework upper estimate and an independently constructed lower one, and a sequence-level check corroborates it only preliminarily. This independently reproduces, with a partition-free instrument and at the monocot—eudicot distance, the classic observation that cross-module connectors are more conserved than hubs whose links stay within a single module [Guimerá and Amaral, 2005]—a finding originally obtained on metabolic networks through the participation-based role cartography. That a measure requiring no partition recovers the same conservation signal on a different kingdom and network type is evidence for the robustness of the underlying biology, not against it.

What this organizational class corresponds to functionally is a question our results open rather than settle. The low-*z*, non-essential connectors EDVS isolates are broadly coordinating yet locally non-dominant, and it is tempting to read them as coordinating loci that might be tuned rather than only broken—including the yield—stress trade-offs that pleiotropy imposes on crop improvement [Paaby and Rockman, 2013]. Our pre-registered probes for such a phenotypic correlate (essentiality, transcription-factor identity, tissue-specificity, date/party-hub character, and yield—stress QTL co-association) found none at the resolution current public references afford (Section 2.7); we therefore present this reading as an open hypothesis the method makes testable, not as a result. The more immediate question is mechanistic: *why* genes come to occupy this position—what evolutionary and functional processes generate the cross-module, locally non-dominant connectivity the measure detects. Mechanism 1 fixes the topological side of that question in direction; sharpening it into a closed-form bound and identifying its biological counterpart is a well-posed program the class now makes addressable with a single annotation-free instrument.

These conclusions carry defined bounds, and one comparison in particular we are explicit about not making: *we do not claim that EDVS is the most accurate classifier of Guimerá-Amaral role labels*. On synthetic networks with planted connector labels, partition-based methods score highest— but they do so using a community partition that, re-detected, nearly recovers the planted one, so that the comparison scores agreement with the label’s own definition rather than recovery of structure out of sample (full benchmark in the Supplementary Material). EDVS does not compete inside that definition; its claim is the different, partition-free one—that it reaches the connector region of the *z*-*P* plane, at its low-*z* extreme, without computing or choosing any partition, and holds that position when the partition choice, and with it the label, moves. This scope is a choice, not an omission: the audit’s target is the criterion in practical use.

The value of a partition-free instrument is sharpest exactly where community detection is least trustworthy. In real biological networks—sparse, noisy, and pervaded by false-positive edges—a reliable community partition is a luxury rather than a given, and the participation-based cartography that defines the connector role inherits that fragility: the very coordinates in which a non-hub connector is defined require a partition the data may not support. EDVS answers this not by improving the partition but by removing the need for one, reaching the low-*z* connector region from a more basic quantity—the entropy of local neighbourhood connectivity—under fewer assumptions and with greater stability to perturbation. EDVS is not the best connector classifier in the abstract: where a trustworthy partition is available the participation coefficient remains the direct instrument. But for the non-hub connector, in the regime where partitions are unreliable, EDVS is among the most suitable selectors available without one—the purest in its low-*z* isolation and the most robust to edge noise, comparable to a spectral removal criterion on the synthetic benchmark, ahead in some regimes and behind in others rather than uniformly best (Supplementary Material) —and that regime is the common case, not the exception. It does not replace participation where a partition can be trusted; it fills the gap where one cannot.

None of this displaces the measures already in use, and we do not read it as a contest to be won. Degree surfaces the provincial hubs and locally important genes it is built to surface; betweenness surfaces the connector hubs that bridge modules while remaining locally central; EDVS supplies the low-*z* non-hub connector that both omit. The practical recommendation that follows is not to replace degree or betweenness but to read a network through the several measures together, with EDVS added as the instrument that recovers a region the standard tools leave dark. Within these bounds, EDVS is an annotation-free, partition-free instrument that isolates a distinct connector class defined by organizational position, requiring neither a community partition nor all-pairs shortest paths. It is not, we are careful to say, a cheap one: it is faster than betweenness but not than a Leiden partition, and carries the largest memory footprint of the three (Section 4.2). And it brings a measurement-theoretic discipline—evaluation against a functional-diversity reference—to the study of network centrality.

## 4 Methods

### 4.1 Networks and functional annotations

The primary organism is rice (*Oryza sativa*) and the primary network is the co-functional network RiceNet v2 [Lee et al., 2015a]. Because RiceNet edges carry a wide range of confidence (log-likelihood scores, LLS, from 1 to 5) and the network’s authors report substantially higher precision for high-scoring links, we take as the canonical network the high-confidence subset (LLS ≥ 1.9519; the top 588,221 edges, ~33%, the threshold adopted by the RiceNet authors; *N* = 20,851 genes). All analyses are repeated on a less stringent ~50% subset and on the full unfiltered network (LLS *>* 0) as sensitivity analyses, and quantities that shift with the threshold are reported explicitly. For within-species cross-network comparison we use two *Arabidopsis thaliana* networks chosen for opposite, deliberate reasons: the co-expression network ATTED-II [Obayashi et al., 2009] —mutual-rank co-expression relations thresholded at *Z* ≥ 3.0, built by an independent laboratory and, importantly for the audit of Section 2.2, by a construction that uses no functional annotation—serves as an *independent* replication of the within-species analyses; AraNet v2 [Lee et al., 2015b], built by the same log-likelihood functional-association framework as RiceNet v2, serves the cross-species conservation analysis, where same-framework construction is required so that cross-species differences are attributable to evolution rather than to network type. The distinction is not incidental: the log-likelihood frameworks (RiceNet v2, AraNet v2) and STRING weight their edges by benchmarking evidence channels against curated co-annotation, whereas ATTED-II and BioGRID do not, and the audit is reported stratified along exactly that line.

To test whether the connector signature generalises beyond plant co-functional networks—across both kingdom and network genre—we add yeast (*Saccharomyces cerevisiae*) networks of two deliberately different construction types. As a functional-association network directly analogous to RiceNet, we use STRING v12 [Szklarczyk et al., 2023] (organism 4932), taking its high-confidence subset (combined score ≥ 700; largest connected component *N* = 5,716) as the canonical yeast network and its medium-confidence subset (score ≥ 400; *N* = 6,290) as a sensitivity analysis. As a genuinely different network *type*, we use the physical protein-protein-interaction network from BioGRID [Oughtred et al., 2021] (physical interactions only; largest connected component *N* = 6,070). For every network, self-loops (*c*_*ii*_ = 1) are added only to the matrix pair on which EDVS is computed, as its definition requires; degree, Burt’s constraint, betweenness, the Leiden partition and the *z*-*P* map are computed on the plain (self-loop-free) topology, matching the rice convention for every measure except EDVS itself.

Functional annotations are Gene Ontology biological-process (GO-BP) terms, propagated to ancestors and restricted to a minimum depth of two to remove uninformative roots, and KEGG pathway memberships—for rice and *Arabidopsis* from their respective GO/KEGG resources, and for yeast from the SGD GO-BP annotation [Cherry et al., 2012] and KEGG `sce` pathways [Kanehisa and Goto, 2000]. Gene essentiality in yeast uses the genome-complete essential-ORF set from the systematic deletion collection [Giaever et al., 2002] (*n* = 1,110, fully resolved to systematic ORF names); this genome-complete label lets the essentiality analysis be run against a complete label rather than the sparse embryo-essential (*EMB*) set available in *Arabidopsis*. One-to-one rice-*Arabidopsis* orthologues for the conservation analysis are taken from the AraNet/Ensembl Plants orthology tables; best-hit orthologues are used as a sensitivity check.

### 4.2 The EDVS centrality

EDVS (Entropy of Degree-Vector Sums) was introduced as an efficient estimator of Rao-Stirling/DIV diversity that satisfies the variety, balance and disparity requirements while computing directly from observed data, without the joint distribution or mutual information the parent indices require [Zhao and Yang, 2023]. Let *A* be the (weighted) adjacency matrix, so that row *A*[*j*, :] is the connectivity (“degree”) vector of node *j*, and let *C* be the binary connectivity matrix, *c*_*ij*_ ∈ {0, 1}, indicating whether *i* is connected to *j*. EDVS requires self-loops, *c*_*ii*_ = 1; networks lacking them are preprocessed to add them. The measure is

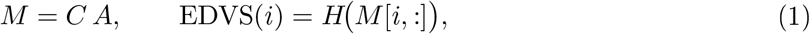

where *H*(·) is the Shannon entropy of the (normalised) row. Equivalently, writing *X*_*j*_ = *A*[*j*, :] and summing over the closed neighbourhood *N* [*i*] = {*j* : *c*_*ij*_ = 1} (which includes *i* through its self-loop),

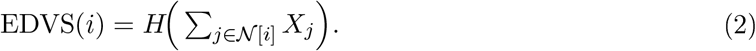

Thus *M* [*i*, :] is the sum of the degree vectors of *i*’s neighbours (and of *i* itself), and EDVS is the entropy of that aggregate: it is large when a node’s neighbourhood connectivity is spread evenly over many, mutually dissimilar targets, and small when it concentrates on a few similar ones.

The original network is a directed, weighted citation network. Our gene networks are undirected, so *A* is symmetric and *N* [*i*] is the ordinary closed neighbourhood; we report EDVS both with *A* binary (EDVS_binary_, the primary score) and with *A* weighted (EDVS_weighted_, a robustness variant). Co-functional and interaction networks have no self-loops, so we add them (*c*_*ii*_ = 1) as the definition requires. Under the binary undirected form, *M* [*i, k*] = ∑ _*j*∈*N* [*i*]_ *A*[*j, k*] counts how many members of *i*’s closed neighbourhood connect to *k*, i.e. the length-two walk mass from *i* to *k*; EDVS is the entropy of this distribution over *k*.

The original formulation has time complexity *O*(*n*^2^), an order below the *O*(*n*^3^) of the Rao— Stirling and DIV indices it approximates [Zhao and Yang, 2023]. On a sparse network the entropy-of-sum form (Eq. 2) computes each node’s aggregate by summing the adjacency rows of its closed neighbourhood, so the cost scales with the number of length-two walks, 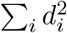, well below the dense *O*(*n*^2^) bound but above 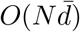 for the heavy-tailed degree distributions of biological networks. What matters for our comparison is categorical rather than constant-factor: EDVS requires neither the all-pairs shortest paths of betweenness (*O*(*nE*), [Brandes, 2001]) nor the community detection that the participation coefficient presupposes, computing the connector signal from local neighbourhood aggregation alone. We stress that this is a structural claim and not an efficiency claim, because measured cost does not support the latter. On our networks EDVS runs 20-80× faster than betweenness (on the 20,851-node rice network, 44.7 s against 1,560 s), but it is not faster than computing a Leiden partition and the participation coefficient from it (19.3 s on the same network), and its peak memory is the highest of the three—5.8 GB against betweenness’s 305 MB—because the *CA* product densifies intermediate rows on networks of this average degree. A chunked implementation would reduce the footprint; we report the naive one we used. Nothing in what follows rests on EDVS being cheap, only on its not requiring a partition.

### 4.3 Why EDVS selects low-within-module-degree connectors

The distinctive behaviour of EDVS—selecting connectors of high participation but low within-module degree—follows from its construction. We make this precise. Let the network carry a community partition with the defining property of community structure: nodes are denser within modules than between them, so that two nodes in the same module share more of their neighbourhoods than two nodes in different modules. For node *i*, the Guimerá-Amaral within-module degree *z*-score *z*_*i*_ increases with the fraction of *i*’s edges that stay inside *i*’s own module, and is maximal when *i*’s connectivity concentrates in one module.

#### Mechanism 1 (heuristic argument, not a theorem)

*Under community structure*; EDVS(*i*) *decreases with the within-module concentration of i s neighbourhood; hence selecting nodes of high EDVS selects nodes of low z*. We state this as a directional mechanism rather than a proposition: the argument below fi es the sign of the effect, not its magnitude, and we do not claim a bound. Its empirical counterpart is the *z*-gradient of Section 2.3.

#### Argument

EDVS(*i*) = *H*(*M* [*i*, :]) with *M* [*i*, :] = ∑ _*j*∈*N* [*i*]_ *A*[*j*, :]. Two structural facts control the shape of *M* [*i*, :]. First (*balance*): *H* is maximised by a uniform distribution and is strictly reduced by concentration of mass. If *i*’s edges concentrate within a single module (high *z*_*i*_), then, by community structure, *i*’s neighbours also lie in and around that module and their degree vectors *A*[*j*, :] place their mass on the same cohesive set of nodes; the sum *M* [*i*, :] is therefore peaked over that small set and *H*(*M* [*i*, :]) is low. If instead *i* bridges several modules (low *z*_*i*_, high participation), the *A*[*j*, :] place mass on disjoint node sets across modules, *M* [*i*, :] spreads over a large support, and *H* is high. Second (*disparity*): summation interacts with neighbour similarity. Within-module neighbours are similar—their degree vectors overlap heavily—so summing them piles mass on shared targets, sharpening the peak and further lowering *H*; cross-module neighbours are dissimilar, so their degree vectors add in near-disjoint coordinates, flattening *M* [*i*, :] and raising *H*. Balance and disparity thus act in the same direction: both penalise the within-module concentration that produces high *z*, so EDVS is structurally anti-correlated with *z*_*i*_.

This mechanism separates EDVS from the reference measures. Degree, deg(*i*) = ∑ _*k*_ *A*[*i, k*], is the total mass of *i*’s own row and is blind to how that mass distributes across modules, so a node with all its edges inside one module (high *z*) can have high degree; degree therefore does not penalise within-module concentration. Betweenness rewards lying on shortest paths; in dense modular networks the inter-module shortest paths funnel through nodes that are also well embedded within a module, so betweenness co-selects higher-*z* connector hubs. Only EDVS takes as its objective the entropy of the aggregated neighbourhood distribution, which is uniquely maximised by cross-module spread, i.e. by low *z*. Mechanism 1 establishes the *direction* of the effect; its magnitude and robustness are given by the empirical *z*-gradient of Section 2.3 (EDVS below betweenness in every network-by-resolution comparison, and lowest of the four arms in seven of nine, now confirmed across kingdoms and network genres, including yeast), which we establish empirically rather than analytically. Sharpening this direction-only argument into a closed-form bound, and asking what biological processes generate the low-*z* connectivity the measure detects, is one of the open questions this work leaves for future study.

### 4.4 Reference centralities and the module map

We compare EDVS against three reference centralities. *Degree* is the node degree. *Burt’s constraint* [Burt, 1992, 2004] measures structural-hole spanning (low constraint indicates a broker across otherwise disconnected contacts); we report its top-1% by lowest constraint. *Betweenness* is computed with Brandes’ algorithm (*O*(*nE*), [Brandes, 2001]). To place all measures on the same organizational map we partition each network with the Leiden algorithm [Traag et al., 2019] at resolutions 0.5, 1.0 and 2.0, and for every node compute the Guimerá —Amaral within-module degree *z*-score and participation coefficient *P [*Guimerá and Amaral, 2005]; *z* measures local dominance within a node’s module and *P* measures how a node’s edges distribute across modules. Throughout, we use “provincial-hub-type” and “connector-type” descriptively, for the higher-*z/*lower-*P* and lower-*z/*higher-*P* ends of the gradient a measure’s selections occupy, and *not* as the formal Guimerá-Amaral role assignments: those rest on absolute cutoffs (*z* ≥ 2.5) whose membership is itself strongly resolution-dependent (on the canonical rice network the degree-exclusive set is 100% above the hub cutoff at the coarsest resolution but 14-16% at finer ones, as modular *z* compresses when modules shrink), so we do not rely on them to classify measures.

### 4.5 Exclusive sets

Several analyses contrast the genes *uniquely* selected by each measure: its top-1% set minus the top-1% sets of the others (its “exclusive” set). Where betweenness is itself under comparison (the *z*-*P* map, edge-architecture and diversity analyses) the e clusion is four-way (degree, Burt, betweenness, EDVS), giving on the canonical rice network EDVS-exclusive *n* = 150, betweenness 82, degree 69, Burt 32; the same four-way construction is used in yeast, with a pre-registered top-2% sensitivity because yeast networks are smaller and small exclusive sets are less stable. The same four-way definition is used in the annotation- and phenotype-based analyses (transcription factors, essentiality, funRiceGenes), so that every exclusive-set analysis rests on a common definition; the canonical EDVS-exclusive set is thus *n* = 150 throughout. For cross-species conservation the focal set is further restricted to genes with a 1:1 orthologue in the target network (*n* = 125; the protein-identity comparison uses the *n* = 52 of these with identity data). Set sizes therefore differ between analyses and are stated wherever used.

### 4.6 Functional-diversity evaluation

For a selected gene set we compute its functional diversity as the rarefied Shannon entropy of the multiset of propagated GO-BP terms of its members, rarefaction controlling for set size. The reference is an annotation-count-matched maximum-entropy expectation: random gene sets drawn to match the selected set’s per-gene annotation counts, which fi es the “research-attention” confound (better-studied genes carry more terms) and asks only whether a set’s functions are concentrated or spread relative to that matched null. We test each measure’s set against this reference by two one-sided tests (TOST) for equivalence within a margin of ±0.15 bit, and report bootstrap 90% confidence intervals. The margin was fi ed in advance at ±0.15 bit, chosen on grounds independent of the results: it is of the order of the bootstrap uncertainty on a single set’s rarefied entropy, and about 2% of the reference entropy itself, so that “equivalent” cannot be attained merely by an underpowered test. *Post hoc*, it proved to be 5-20× smaller than the collapse effects it must not absorb (degree- and Burt-selected sets fall 1.0-3.3 bit below the reference); a set is “collapsed” if significantly below the reference and “preserved” if equivalent to or above it. We additionally report the number of distinct pathways covered by each set (set-level coverage), which unlike per-gene term counts is a property of the set rather than of its individual genes.

### 4.7 Coverage-mechanism analyses

To ask what drives the functional coverage a selection attains (Section 2.5), we fit a partial-effects O S regression of coverage (both the rarefied-entropy difference and set-distinct-pathway count) on within-module degree *z* or participation *P* together with mean degree, separately per axis and per network-resolution combination (15 combinations: 5 networks × 3 resolutions), to separate each axis’s effect from the degree confound its selections share. A non-parametric cross-check bins genes into degree quintiles (5 bins, minimum 40 genes per bin) and compares coverage between matched high- and low-purity subsets within each bin, using 150 subsampling replicates; agreement with the regression is not assumed and disagreement is reported rather than resolved by preferring one method. Whether participation’s effect on coverage runs through the number of distinct modules a selection spans is tested with a Sobel mediation test on the *P* → modules-spanned → coverage chain, controlling for degree throughout. Partition-choice sensitivity (Section 2.5) is assessed by recomputing the top-*P* selection under nine partitions: Leiden and Louvain [Blondel et al., 2008], each at resolutions 0.5, 1.0 and 2.0, and Infomap [Rosvall and Bergstrom, 2008] under three random seeds (Infomap has no resolution parameter in our implementation, so seed rather than resolution is varied). Edge-perturbation robustness is assessed two ways: selection stability under random 10% edge deletion, with Jaccard between the perturbed and unperturbed top-1% sets over 5 (rice) or 10 (yeast) replicates; and coverage stability under combined edge deletion and injection at 10%, 20% and 30% of the original edge count (one replicate per level). All procedures use seed 42.

### 4.8 Enrichment and permutation nulls

Enrichment of an exclusive set for an e ternal label—transcription factors (Plant TFDB), embryo-essential *EMB* genes in *Arabidopsis* (SeedGenes), functionally characterised rice genes (funRiceGenes), or essential ORFs in yeast (the systematic deletion collection, Giaever et al., 2002) —is tested against a *degree-matched* permutation null: for each focal gene we draw, from a pool binned by degree, a matched non-focal gene, and repeat ≥ 2000 times to obtain the null distribution of the label proportion, reporting the observed proportion, its *z* and empirical *p* against that null, alongside the raw Fisher test. Degree matching is essential because all top-1% selections are degree-elevated; only the degree-matched null asks whether a label is enriched *beyond* what degree alone e plains—a distinction that matters especially for yeast essentiality, where the classical centrality-lethality correlation makes high-degree selections essentiality-elevated by construction. Effect sizes for continuous contrasts use Cliff’s *δ*. Because the same contrast is repeated across networks, resolutions and set sizes, we treat each such family as a pre-registered replication rather than a search: the hypothesis, direction and decision rule were fixed before results were inspected, and we report every cell of each family—including null and reversed ones—rather than the best. Where a family is exploratory rather than pre-registered we say so explicitly and apply Benjamini-Hochberg control within it. Pre-registration constrains the choice of test, not the number of cells, so replication counts (e.g. 9/9, 13/15) should be read as consistency across conditions rather than as independent confirmations.

### 4.9 Probes for a phenotypic or importance signal

To ask whether the organizational connector class EDVS isolates also carries a detectable phenotypic or importance signal, we ran a set of pre-registered probes against external references, each with the degree-matched null above: gene essentiality (yeast systematic-deletion set; *Arabidopsis EMB*); transcription-factor identity; tissue-specificity of expression (the *τ* index over a rice expression compendium); date-hub versus party-hub character (mean co-expression of a gene with its network neighbours over an independent rice RNA-seq compendium); and co-localisation with, and trade-off co-association across, rice yield and abiotic-stress QTLs (Q-TARO trait categories). These probes are reported for scope (Section 2.7); full pre-registered protocols, data sources and per-probe statistics are given in the Supplementary Material. Each was frozen (hypothesis, direction and analysis fixed) before results were inspected, and null or reversed outcomes are reported as such. Because a non-significant test is not an absence, each probe is additionally assessed for *equivalence* to its degree-matched null by two one-sided tests within a margin of ±5 percentage points on the label proportion, with 90% intervals—the same procedure as the functional-coverage evaluation above. This margin was set for the binary labels after the probes had been run, so we report it as a post-hoc equivalence assessment rather than a pre-registered one, give the interval half-widths so that sensitivity to the margin can be judged directly, and distinguish equivalence from inconclusiveness wherever an interval exceeds the margin.

### 4.10 Cross-species conservation

To test whether the connector position is conserved, we map rice EDVS-exclusive connectors to their 1:1 *Arabidopsis* orthologues and compare the orthologues’ EDVS percentile in the target network against a rice-degree-matched null, reporting the percentile shift, its *z* and empirical *p*, in both directions (rice→ath and ath→rice). Degree-independence is assessed by a paired contrast on the same focal genes: the role-specificity Δ = (EDVS-percentile shift) − (degree-percentile shift), with a 90% bootstrap CI. Because AraNet v2 and RiceNet v2 share a construction framework, the AraNet effect is treated as an *upper* bound that cannot fully exclude shared-construction bias, and the independently constructed ATTED-II provides a method-independent *lower* bound. A sequence-level check uses percent protein identity of 1:1 orthologues (dN/dS being unavailable at the monocot-eudicot distance owing to synonymous-site saturation); it is reported as preliminary.

